# SYNAPTIC RELEASE POTENTIATION AT AGING AUDITORY RIBBON SYNAPSES

**DOI:** 10.1101/2020.05.15.097550

**Authors:** Thibault Peineau, Séverin Belleudy, Susanna Pietropaolo, Yohan Bouleau, Didier Dulon

**Affiliations:** Neurophysiologie de la Synapse Auditive, INSERM UMRS 1120, Bordeaux Neurocampus, Université de Bordeaux, 33076 Bordeaux, France; Institut de l’Audition, Centre Institut Pasteur/Inserm, 63 rue de Charenton, 75012 Paris; INCIA, UMR 5287, CNRS, University of Bordeaux, Bat B2, Allée Geoffroy St. Hilaire, CS 50023, 33615, Pessac Cedex, France

**Keywords:** Auditory hair cells, Synaptic ribbons, Synaptopathy, Hyperacusis, Ca^2+^ channels, Exocytosis

## Abstract

Age-related hidden hearing loss is often described as a cochlear synaptopathy that results from a progressive degeneration of the inner hair cell (IHC) ribbon synapses. The functional changes occurring at these synapses during aging are not fully understood. Here, we characterized this aging process in IHCs of C57BL/6J mice, a strain which is known to carry a cadherin23 mutation and experiences early hearing loss with age. These mice, while displaying a large increase in auditory brainstem thresholds due to 50 % loss of IHC synaptic ribbons at middle age (postnatal day 365), paradoxically showed enhanced acoustic startle reflex suggesting a hyperacusis-like response. The auditory defect was associated with a large shrinkage of the IHCs’ cell body and a drastic enlargement of their remaining presynaptic ribbons which were facing enlarged postsynaptic AMPAR clusters. Presynaptic Ca^2+^ microdomains and the capacity of IHCs to sustain high rates of exocytosis were largely increased, while on the contrary the expression of the fast repolarizing BK channels, known to negatively control transmitter release, was decreased. This age-related synaptic plasticity in IHCs suggested a functional potentiation of synaptic transmission at the surviving synapses, a process that could partially compensate the decrease in synapse number and underlie hyperacusis.

## INTRODUCTION

Age-related hearing loss (ARHL or presbycusis) is the most prevalent form of sensory disability in human populations and adversely affects the quality of life of many elderly individuals. Simultaneously with ARHL, chronic tinnitus and hyperacusis often develop due to abnormal neural activity in the central auditory system whose origin may arise from cochlear pathologies (for review see Knipper et al., 2013). The development of ARHL has been generally considered to be associated with primary degeneration of cochlear hair cells, with secondary deafferentation associating degeneration of spiral ganglion neurons and ribbon synapses. However, recent studies have resulted in a paradigm shift in the understanding of the early cause of ARHL (Kujawa and Liberman, 2015; 2019). During aging, hearing thresholds in the elderly often remain normal but speech intelligibility become hampered, especially in noisy environment. This pathology in the elderly is referred as hidden hearing loss and has been proposed to be caused in mice by an early loss of ribbon synapses contacting the inner hair cells (IHCs), well before any major loss of hair cells and spiral ganglion neurons (Stamataki et al., 2006). While glutamate excitotoxicity, resulting in nerve terminal swelling, has been proposed to play a role in the setting of noise-induced hearing loss (Pujol and Puel, 1999; Hu et al., 2020), the molecular mechanisms underlying the degeneration of the cochlear ribbon synapses during aging remain to be elucidated. Oxidative damage due to an increase in ROS levels seems to plays a crucial role in ARHL (Fu et al., 2018) but how the ribbon synapses are functionally affected remain unknown.

The C57BL/6J inbred strain of mice has been extensively used as model for ARHL (Johnson et al., 1997; Spongr et al., 1997; Hequembourg and Liberman, 2001; Bartolomé et al., 2002; Henry, 2002). This mouse strain carries a cadherin23 mutation (*Cdh23* ^753A^, also known as *Ahl*), which affects inner ear structures, characterized by progressive degeneration of ribbon synapses and spiral ganglion neurons, and results in age-related hearing loss. *Ahl* is a recessive single nucleotide mutation at 753 (G=>A) on the *Cdh23* gene on mouse chromosome 10 (Noben-Trauth et al., 2003). *Cdh23* encodes cadherin 23, a protein necessary for inner ear development and maintenance of the sensory hair cell structures, in particular the tip and lateral links of the stereocilia at the hair bundle (Siemens et al., 2004; Söllner et al., 2004; Kazmierczak et al., 2007; Michel et al., 2005; Lagziel et al., 2005), but growing anatomical evidence also suggests that synaptic rearrangements on sensory hair cells also contribute to auditory functional decline in C57BL/6J mice (Stamataki et al., 2006; Zachary and Fuchs, 2015; Jiang et al., 2015; Jeng et al., 2021). Surprisingly, aging C57BL/6J mice, although showing cochlear ribbon synapse degeneration and hearing loss, display increased acoustic startle reflex amplitudes (Ouagazzal et al., 2006; Ison and Allen, 2003; Ison et al., 2007) and a time increase in recovery from ABR short-term adaptation (Walton et al., 1995). Whether this exaggerated startle reflex can be explained by functional changes of cochlear hair cell ribbon synapses remain to be explored. Our present study will show that aged C57BL/6J mice, while losing nearly up to 50 % of their IHC ribbon synapses, have IHCs with reduced BK channel clusters, larger remaining presynaptic ribbons (with increased Ca^2+^ microdomains and larger sustained exocytotic responses) facing postsynaptic afferent boutons with larger AMPAR clusters. All these features indicated that synaptic release potentiation occur at auditory IHC ribbon synapses during aging. This age-related synaptic plasticity could be viewed as compensatory mechanisms of the decrease IHC synapse number and may explain the hyperacusis-like startle reflex observed in aged C57BL/6J mice.

## RESULTS

### Increased ABR thresholds and hyperacusis-like effect on ASR

As previously described (Zheng et al., 1999; Zhou et al., 2006), CBA/J mice, used here as a reference, showed no sign of rapid degradation of their hearing function with aging, as attested by constant click ABR thresholds near 15 dB SPL up to one year of age (Fig.1A). On the contrary, C57BL/6J mice, carrying the *Cdh23*^753A^ mutation, displayed an early progressive increase in ABR thresholds with aging (Fig.1A). Hearing thresholds started to increase significantly at postnatal day P180 to reach a mean 50 dB increase at P365 (click ABR thresholds above 65 dB SPL) as compared to young mature P30 mice. Distortion product otoacoustic emissions (DPOAEs) in P365 C57BL/6J mice were still present but significantly reduced, indicating that the electromechanical activity of the outer hair cells (OHCs) was also affected (Fig.1D). However, counting hair cells on surface preparations of organs of Corti from these P365 mice, in the mid-cochlea region (8-16 kHz), indicated only a moderate loss of OHCs and IHCs (statistically not significant; Fig.2E), in good agreement with the study of Zachary and Fuchs (2015). This moderate loss of OHCs suggested that the profound hearing loss occurring at P365 in C57BL/6J mice arose from additional factors than a simple alteration of the cochlear amplifier. We then compared the amplitude on the ABR wave-I as a function of sound intensity in young and old mice. The wave-I is known to result from the synchronous activity of the auditory nerve fibers and can provide an objective measure of ribbon synapses loss when measured at high sound intensity above 70 dB SPL. In P365 old mice, ABR wave-I amplitudes were largely reduced as compared to young mature P30 mice, even at high sound intensity when IHCs are directly stimulated and bypass the amplification process of the OHCs (Fig.1B). The large decrease in wave-I amplitude indicated a decreased number of auditory nerve fibers activated by sound and/or a decrease in their firing rate or synchrony. The latencies of the ABR wave-I were also significantly increased suggesting again that the IHC ribbon synapses are affected (Fig.1C).

Surprisingly, at lower sound intensities tested (between 20-35 dB above background noise level), the strength of the acoustic startle reflex (ASR) was found larger in old P365 C57BL/6J mice as compared to young P60, indicating hyperacusis reactivity (Fig 1.E). The origin of this acoustic hypereactivity at low sound intensities could arise from central neural reorganization (Willott 1996) and be consecutive to functional defect of the IHC ribbon synapses in aging C57BL/6J mice that we are going to characterize in this study. At loud sound intensities, 45 dB above background noise, the amplitude of the ASR responses were however diminished in old mice, likely reflecting the increase in ABR thresholds due to a drastic decrease in the number of ribbon synapses per IHCs.

**Figure 1.**
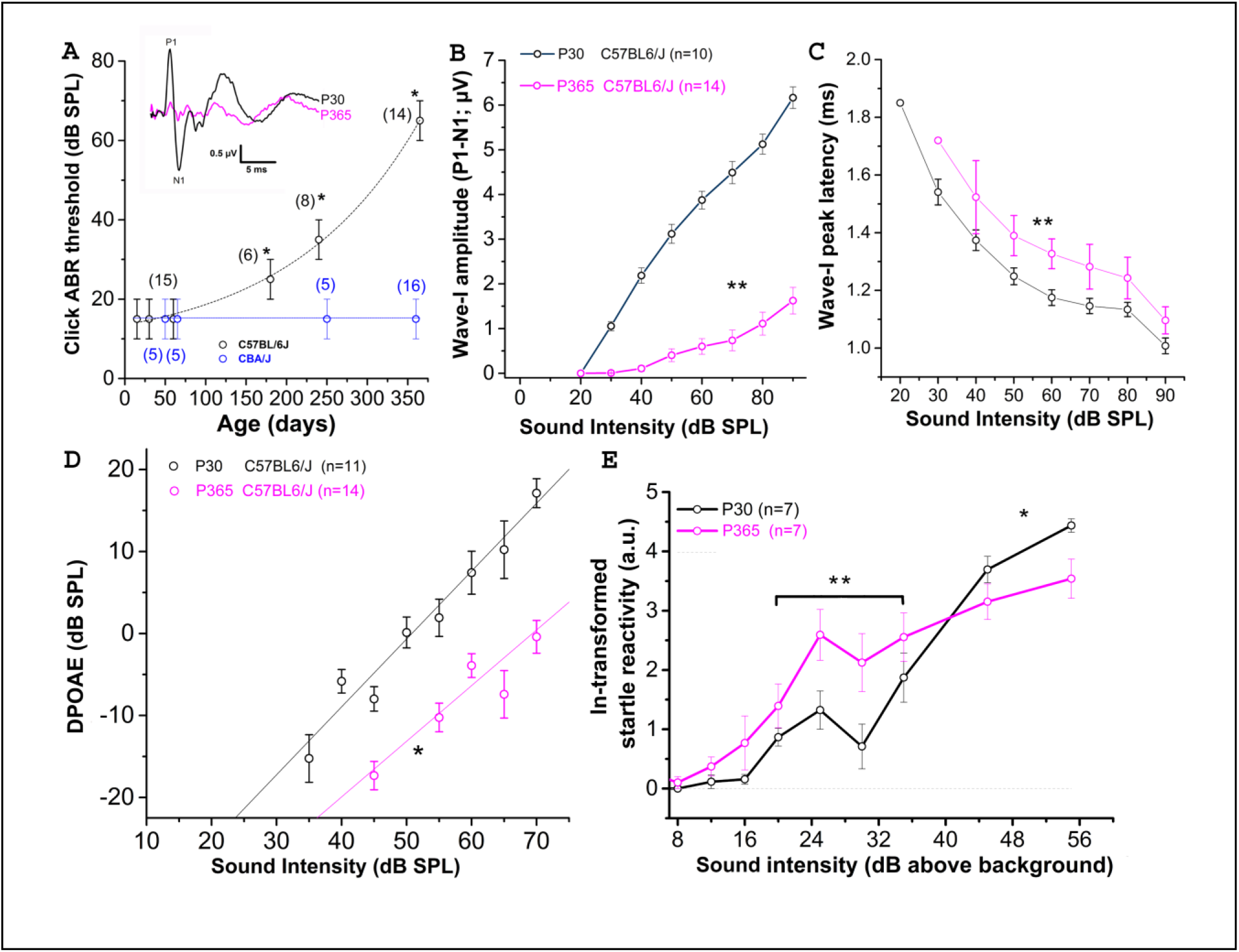
Sensorineural hearing loss with Aging in C57BL/6J mice. (**A**) Click-ABR thresholds as a function of age in CBA/J mice (blue; n = 5 to 16) and C57BL/6J mice (black, n = 6 to 15). For each strain, asterisks indicate significant difference with p<0.05 as compared to P30 mice (two-way ANOVA). Inset show comparative mean ABR waves at 70 dB SPL in P30 and P365 C57BL/6J mice. (**B**) Wave-I amplitude (P1-N1) as a function of sound intensity in young P30 (black) and old P365 (pink) C57BL/6J mice. Asterisks indicate significant difference with p<0.05 (two-way ANOVA; p = 0.001). (**C**) Latency to reach peak amplitude of wave-I as a function of sound intensity. Significantly different with p=0.003 between 80 to 50 dB SPL (two-way ANOVA; n is indicated in graph B). (**D**) Distortion products otoacoustic emissions (DPOAEs; 2f1-f2 with f1 = 12.73 kHz and f2 = 15,26 kHz) as a function of sound intensity in young P30 (black) and old P360 (pink) C57BL/6J mice. Asterisk indicated significant difference with p<0.05 (two-way ANOVA, p = 0.01). (**E**) Startle reactivity to acoustic stimulation. The intensity of startle reaction (in mV) was ln-transformed and obtained through two conditions of white-noise bursts of 20 and 40 ms. Note that below 35 dB above background the startle reaction is increased at P360 whereas at 45 and 55 dB above background the startle reaction is decreased compared to P30 mice. Asteriks indicated significant difference with p<0.05 (two-way ANOVA, p = 0.007 between 20-35 dB and p = 0.005 between 45-55 dB).

### Drastic loss of IHC synaptic ribbons with aging

Confocal immunofluorescence imaging of the synaptic ribbons in IHCs from C57BL/6J mice, in the cochlear region encoding between 8-16 kHz, showed a progressive and large decrease in the number of synaptic ribbons with aging, from 16.3 ± 0.3 ribbons/IHC at P15 to 8.7 ± 0.5 ribbons/IHC at P365 (Fig.2A-C). This nearly 50% loss of synaptic ribbons in IHCs of C57BL/6J mice at P365 was in good agreement with previous studies (Zachary and Fuchs, 2015; Jeng et al., 2020). Counting the spiral ganglion neurons in the mid-frequency region of the cochlea (8-16 kHz) showed a mean 20 % loss of neuron in P365 mice as compared to P30 mice (Fig.2D-F). Remarkably, the age-related loss of neurons, averaging 20 %, was much less pronounced than the loss of IHC synaptic ribbons (50 %), suggesting that there was a large number of neurons surviving without dendritic/synaptic projection toward the organ of Corti as previously shown by Stamataki et al. (2006).

**Figure 2.**
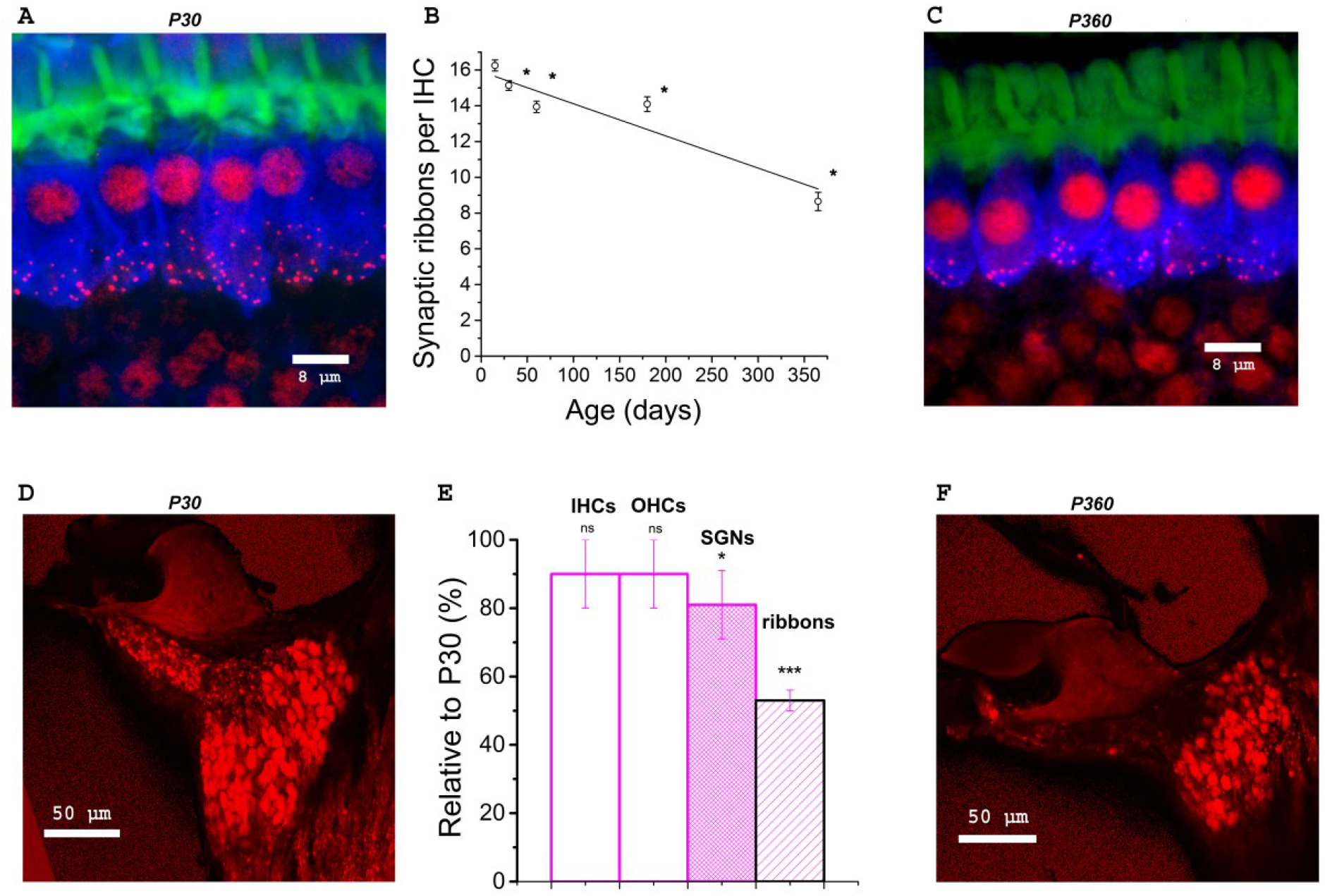
Synaptic ribbon loss in aging IHCs and comparative density of spiral ganglion neurons. **(A-C**) Confocal immunofluorescence imaging of IHCs from P30 and P360 C57BL/6J mice. Image resulted from stack reconstructions of 22 slices of 0.3 µm thickness, showing F-actin (green, labeling the hair bundles), otoferlin (blue; a synaptic Ca^2+^ sensor labeling the entire IHC body) and synaptic ribbons (Ribeye/CtBP2, red dots). The white bars indicated 8 µm. (**B**) Plot of synaptic ribbon counts per IHC as a function of age in C57BL/6J mice (black) in the mid-frequency cochlear region (8-16 kHz). Asterisks indicate significant difference with p<0.05 (comparison with P15 C57BL/6J mice, one-way ANOVA post-hoc Tukey test). 203, 306, 306, 306, 295 ribbons were respectively analyzed at P15, P30, P60, P180, P365 in 3 to 5 mice at each age. (**D-F**) Confocal immunofluorescence imaging of cochlear transversal sections from P30 and P360 mice (8-16 kHz area) where neurons and their afferent dendrites are labeled with dextran amines 3000. (**E**) Analyzed in the mid-cochlea (8-16kHz), comparative histogram of hair cells counts, SGNs and synaptic ribbons (3 mice at each age relative to young mature P30 C57BL/6J mice. The asterisks indicated statistical difference with p < 0.05 (from left to right p = 0.37, p = 0.14, p = 0.020 for SGNs and p = 1.01E-14 for ribbons, using unpaired t-test). Note the reduction in SGNs and the drastic decrease of the fluorescent labeling of the afferent nerve fibers projecting to the organ of Corti.

### Enlarged presynaptic ribbons associated with larger postsynaptic AMPAR clusters

When analyzing the 3-D imaging reconstruction of the IHC synaptic ribbons from P365 C57BL/6J IHCs from high resolution confocal STED immunofluorescent sections, we found that these structures displayed a progressive increase in their mean size (volume) as compared to young P15 mice (Fig.3A). The mean volume of the synaptic ribbons increased nearly three-fold between P15 and P365 (Fig.3B). Interestingly, the increase in size of the ribbons started very early after P30, an age at which their number also started to decrease (Fig.2B) while hearing thresholds were still normal (Fig.1A).

**Figure 3.**
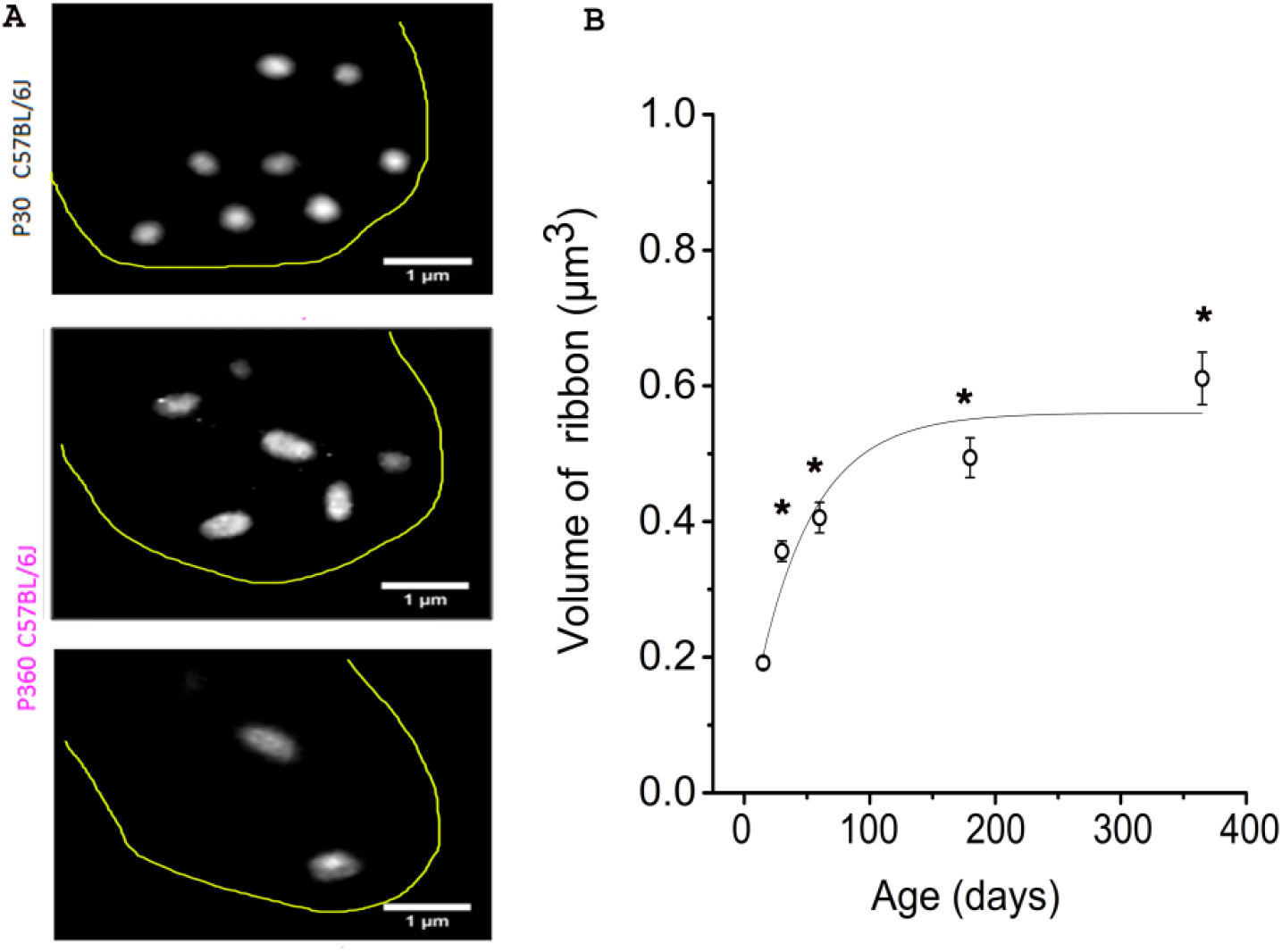
IHC synaptic ribbons progressively increased in size with aging. (**A**) Example of high magnification confocal STED images (stack reconstruction of five 0.25 µm slices). The yellow line delineates the IHC contour. Ribbons were less numerous and larger in size in IHCs from P365 C57BL/6J mice. (**B**) The mean volume of the IHC synaptic ribbons was calculated in the mid-cochlear region (encoding 8-16 kHz), using the plugin 3D-Object counter of ImageJ, and was plotted as a function of age: P15 (203 ribbons -40 IHCs -3 mice), P30 (306 ribbons -98 IHCs -5 mice), P60 (306 ribbons -95 IHCs -3 mice), P180 (306 ribbons -67 IHCs -3 mice), P365 (295 ribbons -43 IHCs -3 mice). As indicated by the asterisks, when comparing with P15, the mean volume of the ribbons started significantly to increase significantly from P30 (p<0.05; one-way ANOVA).

We next investigated the changes occurring with aging in the spatial distribution of the synaptic ribbons at the basolateral membrane of the IHCs. It is worth recalling that larger presynaptic ribbons associated to low-spontaneous-rate afferent fibers are usually found at the modiolar side of the IHCs (i.e. the baso-lateral curved side of the IHCs directed toward the spriral ganglion neuron) while smaller ribbons associated with high-spontaneous-rate afferent fibers predominates at the pillar side (Liberman, 1982; Liberman et al., 2011). We found that, while a larger proportion of synaptic ribbons distributed at the modiolar side in young P30 IHCs (∼60 %), the surviving ribbons in old P365 IHCs were evenly distributed at the pillar and modiolar side (Fig.4A-B). The loss of synaptic ribbons with aging was more pronounced at the modiolar side, ie at the side where most of the low-spontaneous (high threshold) fibers make contact to IHCs (Fig 4C). An increase in volume was noted both in pillar and modiolar ribbons (Fig 4D). Unlike young P30 IHCs, old P365 IHCs had modiolar and pillar ribbons with similar volume (0.7 ± 0.1 µm^3^ and 0.57 ± 0.07 µm^3^, respectively; not significantly different, unpaired t-test p = 0.28, Fig.4D). Similar preferential loss of synaptic modiolar ribbons associated with an increase in ribbon volume was also reported after noise-induced hearing loss (Liberman and Kujawa, 2017; Hickman et al., 2020).

**Figure 4.**
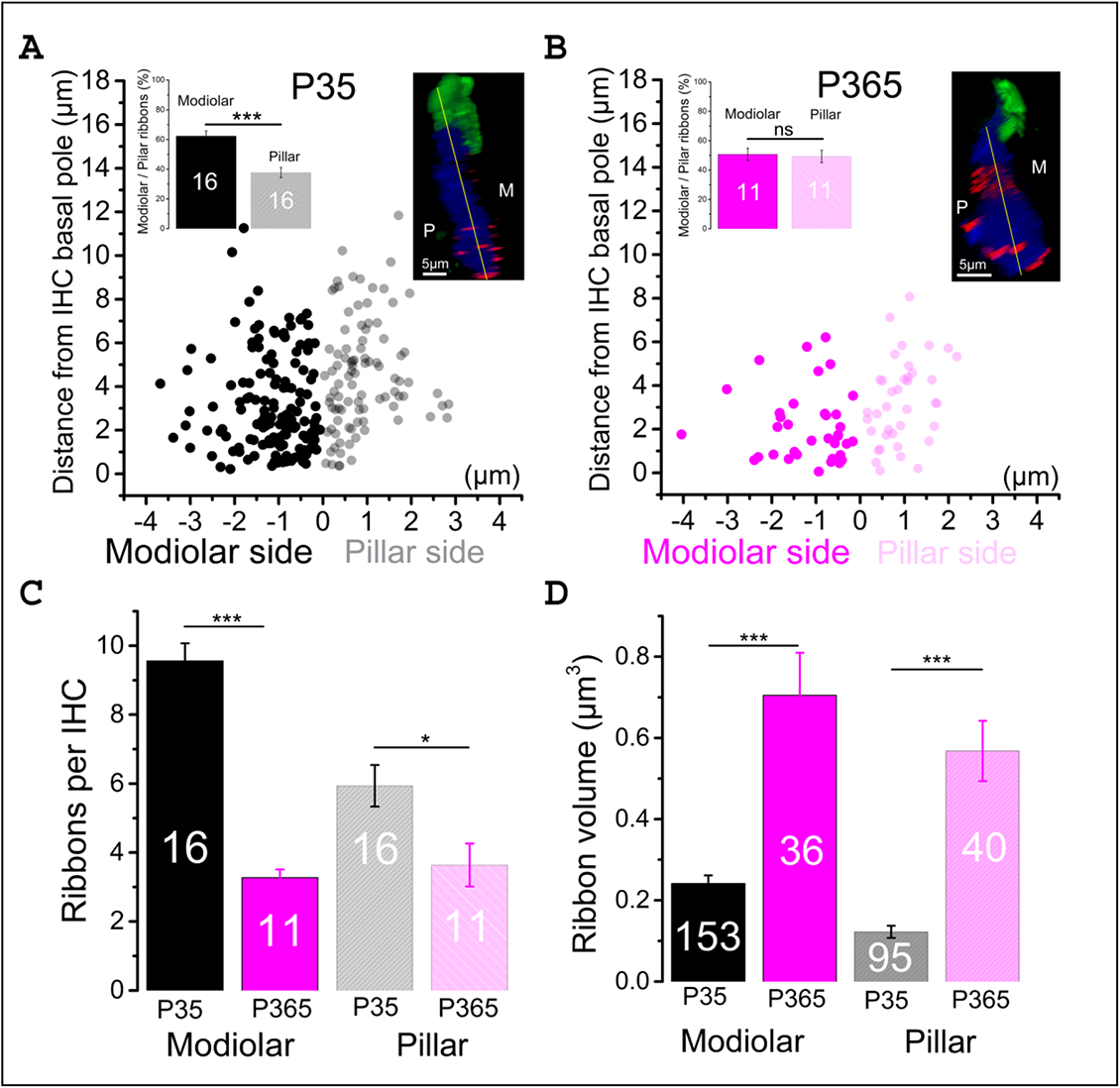
Modiolar vs pillar distribution of the synaptic ribbons with aging. (**A-B**) The distribution of IHC synaptic ribbons was splitted in two cell areas: P (pillar side) and M (Modiolar side). The two regions were determined using the K-means clustering method for each cell (see Methods). The yellow lines in the inset confocal images was drawn to arbitrarily delineate the frontier between pillar and modiolar ribbons (green labeling F-actin, blue Otoferlin and red the synaptic ribbons). Note that in young P30 IHCs (n=16), synaptic ribbons are preferentially localized in the modiolar region (62.3 ± 3.5 % vs 37.7 ± 3.5; unpaired t-test p = 2.07 E-5). In old P365 IHCs (n=11), the remaining ribbons equally distributed in the modiolar and pillar side (50.7 ± 4.0 % vs 49.3 ± 4.0 %, unpaired t-test; p = 0.81). (**C**) Synaptic ribbons were significantly loss both at the modiolar side and pillar side. However, the loss was much more pronounced in the IHCs’ modiolar side (9.56 ± 0.5 ribbons vs 3.27 ± 0.24 ribbons; unpaired t-test p =5.57 E-10) and pillar (5.9 ± 0.6 ribbons vs 3.6 ± 0.6 ribbons, unpaired t-test p = 0.016). (**D**) The increase in volume affected both modiolar (P30=0.24 ± 0.02 µm^3^ vs P365=0.7 ± 0.1 µm^3^ unpaired t-test, p = 3.7 E-11) and pillar (P30= 0.12 ± 0.01 µm^3^ vs P365= 0.57 ± 0.07 µm^3^, unpaired t-test p = 7.5 E-14) synaptic ribbons. This comparative analysis was performed in IHCs of the mid-cochlear region (encoding 8-16 kHz).

Postsynaptic afferent endings to ribbon IHCs express AMPA-type ionotropic glutamate [glutamate receptor A2 (GluA2)]-containing AMPA receptors (Matsubara et al., 1996; Hu et al., 2020). To determine whether the organization of postsynaptic AMPARs facing the IHCs ribbon was modified in old P365 mice we performed GluA2 immunostaining and confocal fluorescence imaging (Fig.5). As compared to young ribbon synapses, old synapses had larger ribbons associated with larger AMPAR clusters, suggesting that the changes occurring with aging are both at the pre and postsynaptic side in good agreement with the EM morphological analysis of Stamataki et al. (2006).

**Figure 5.**
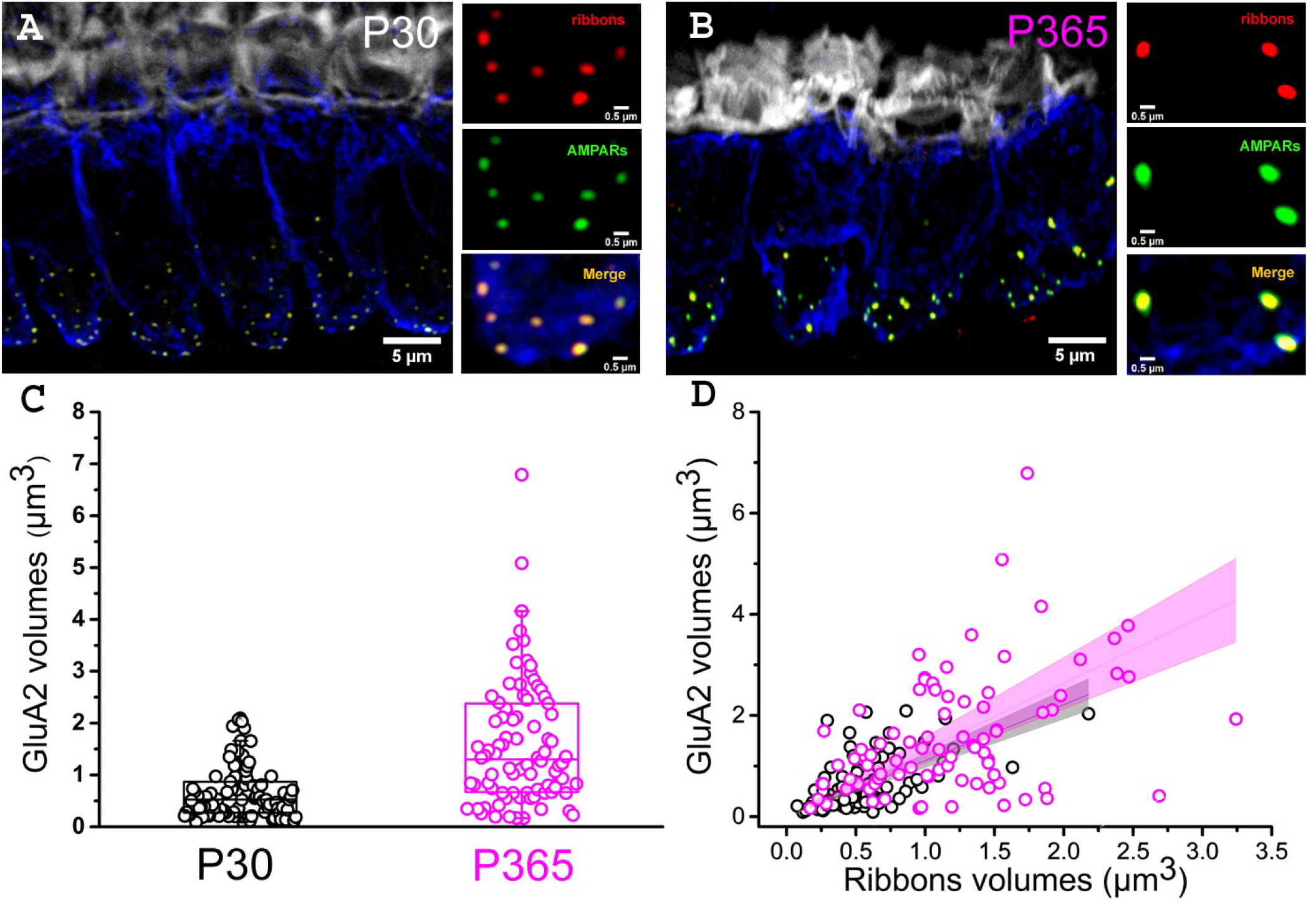
Larger postsynaptic patches of GluA2 receptors in old IHCs. (**A to D**) Immuno-confocal imaging of IHCs (stack reconstructions of 22 slices of 0.3µm thickness) showing otoferlin (blue), the synaptic ribbons (Ribeye/CtBP2, red dots) and the post-synaptic AMPA receptors (GluA2, green dots). Fluorescent-phaloidin (grey) labelled F-actin in the strereocilia The white bars indicate 5 µm in A and B, then 0.5µm in C and D. (**C**) Comparative distribution of the volume of postsynaptic patches of AMPA GluA2 receptors. Note that the GluA2 receptor volume significantly increased with aging in C57Bl/6J mice (p < 0.001 ; unpaired-t test); (n = 86 synapses; from 3 mice at P30 (16 IHCs) and P365 (29 IHCs) ; mid-cochlear region encoding 8-16 kHz). (**D**) Comparative volume distribution of postsynaptic GluA2 and corresponding postsynaptic ribbon in P30 and P365 IHCs. Note a good linear correlation between GluA2 and ribbon volumes (Pearson’s coefficient r = 0.93 and 0.86, respectively for P30 and P360).

### Drastic IHC cell size reduction wih aging

To determine whether the loss of synaptic ribbons was associated with a cell size reduction in old P365 IHCs, we labeled the cytosol of IHCs with an antibody against myo-7a, an unconventional myosin essential for the hair bundle functional integrity and found in the apical sterocilia as well as the cytoplasm of the hair cells (Hasson et al., 1995). We found a drastic reduction of the basal infra-nuclear synaptic zone of the IHCs from an average of 15.7 ± 0.1 µm (n=34) in P21 IHCs to 12.4 ± 0.3 µm (n=50) in old P365 IHCs (unpaired-t test, p = 1.1 E-13), Fig. 6A-C). The cell size reduction of old IHCs was confirmed by another method using whole-cell patch clamp measurement of the IHC resting membrane capacitance in organs of Corti preparation ex-vivo (Fig.6D-E). P365 IHCs displayed a significant smaller resting capacitance, measured at -70 mV, of 7.9 ± 0.4 pF (n=22) as compared to P21 IHCs (10. 8 ± 0.1 pF, n=44, unpaired t-test, p = 3.7 E-10).

**Figure 6.**
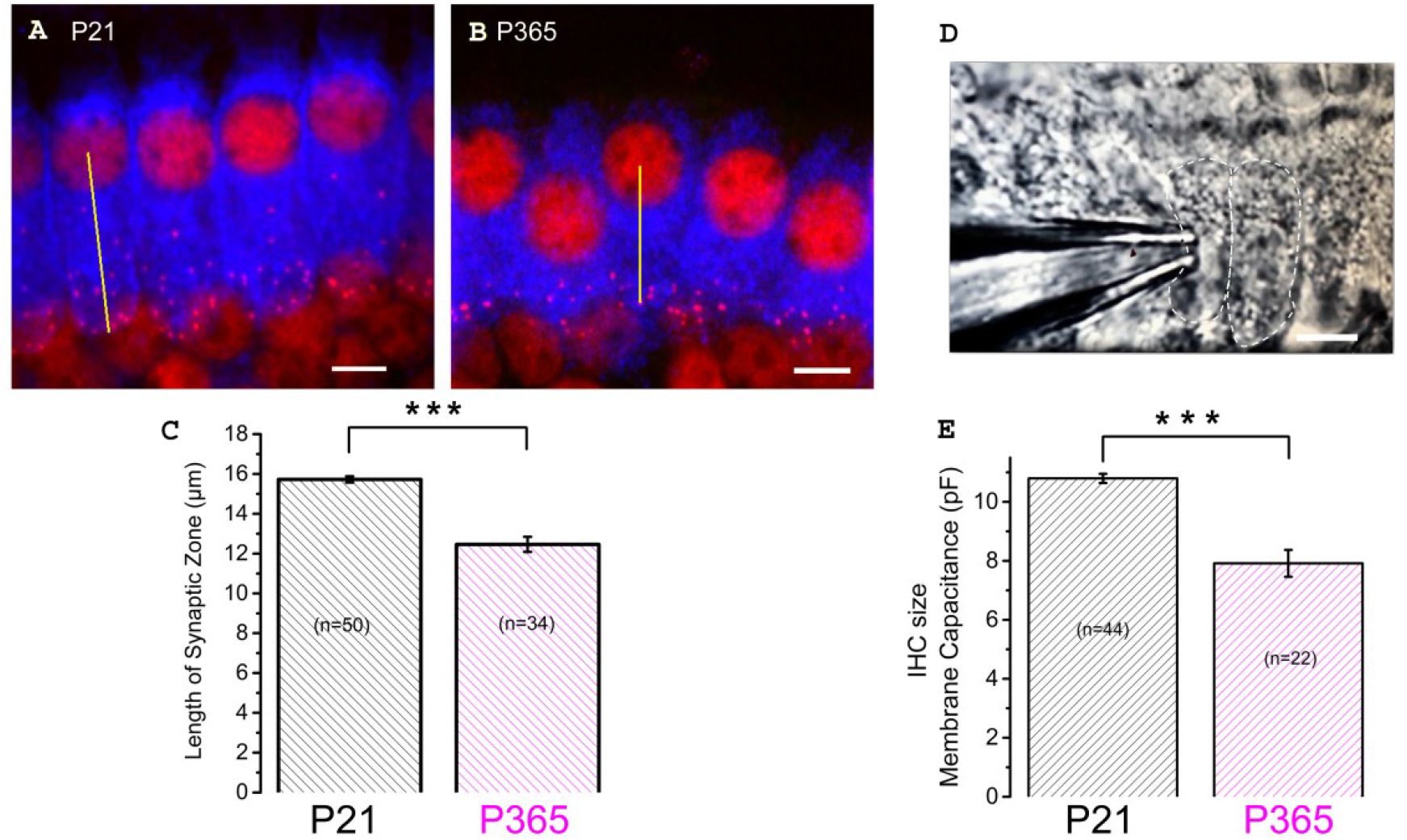
Cell size reduction of IHCs with aging. (A-B) Immuno-confocal imaging of IHCs labeled for myo-7a (blue) and CtBP2 (labeling the ribbons, red) in P21 (A) et P365 (B) mice (reconstructed stack images of 22 slices of 0.3 µm thick). The length of the infra-nuclear synaptic zone was measured from the center of the nucleus to the bottom of the cell as indicated by the yellow line. The scale bar indicated 5 µm. (C) Comparative histogram of the length of the infra-nuclear synaptic zone. Asterisks indicated statistical significance with p < 0.0001, unpaired t-test.) Measurements were made from IHCs of the mid-cochlear region (8-16 kHz). (D) Photomicrograph showing a row of IHCs in an ex-vivo organ of Corti explant and a patch-clamp recording electrode allowing measurements of capacitive and ionic currents. (E) Comparative mean IHC resting membrane capacitance, measured at -70 mV, in whole-cell patch-clamp configuration. Asterisks indicated values with significant difference with p < 0.0001, unpaired-t test.

### Higher Ca^2+^ current density and exocytosis in old IHCs

Although the loss of IHC ribbon synapses in C57BL/6J mice with aging is well documented in the literature (see Wan and Corfas, 2015), the functional changes of the remaining “old” IHC ribbon synapses has not yet been explored in detail at the cellular level. We investigated here the changes in the calcium-evoked exocytotic responses of aging IHCs from ex-vivo explants of the organ of Corti from aging C57BL/6J mice. We recall that Ca^2+^ is a key messenger for normal hearing since it tunes the transduction of the analogic sound signal into nerve impulses. This transduction occurs at the IHC synaptic active zone through the voltage-activation of Ca_v_1.3 Ca^2+^ channels (Brandt et al., 2003; Vincent et al., 2017) and the action of the Ca^2+^ sensor otoferlin (Roux et al., 2006; Beurg et al., 2010; Michalski et al., 2017). Increase in the density of L-type Ca^2+^ channels has long been recognized as a characteristic of aging central neurons (Thibault and Landfield, 1996; Campbell et al., 1996) but possible altered function in Ca^2+^ signaling has not been investigated in aging IHCs.

In the present study, whole cell patch-clamp recordings from old P365 IHCs showed a significantly increase in Ca^2+^ current density (Ca^2+^ current amplitude normalized to cell size) as compared to young mature P30 IHCs, rising from a mean peak value of 15.8 ± 1.75 pA/pF to 20.1 ± 1.9 pA/pF at -10 mV (Fig. 7A; unpaired t-test, p <0.05). The voltage-activation curve of the Ca^2+^ currents in old P365 IHCs remained similar to the young mature P30 IHCs, with a mean half-voltage activation (V_1/2_) of -29.3 ± 1.1 and -26.7 ± 2.8 mV, respectively (unpaired t-test, p = 0.06).

**Figure 7.**
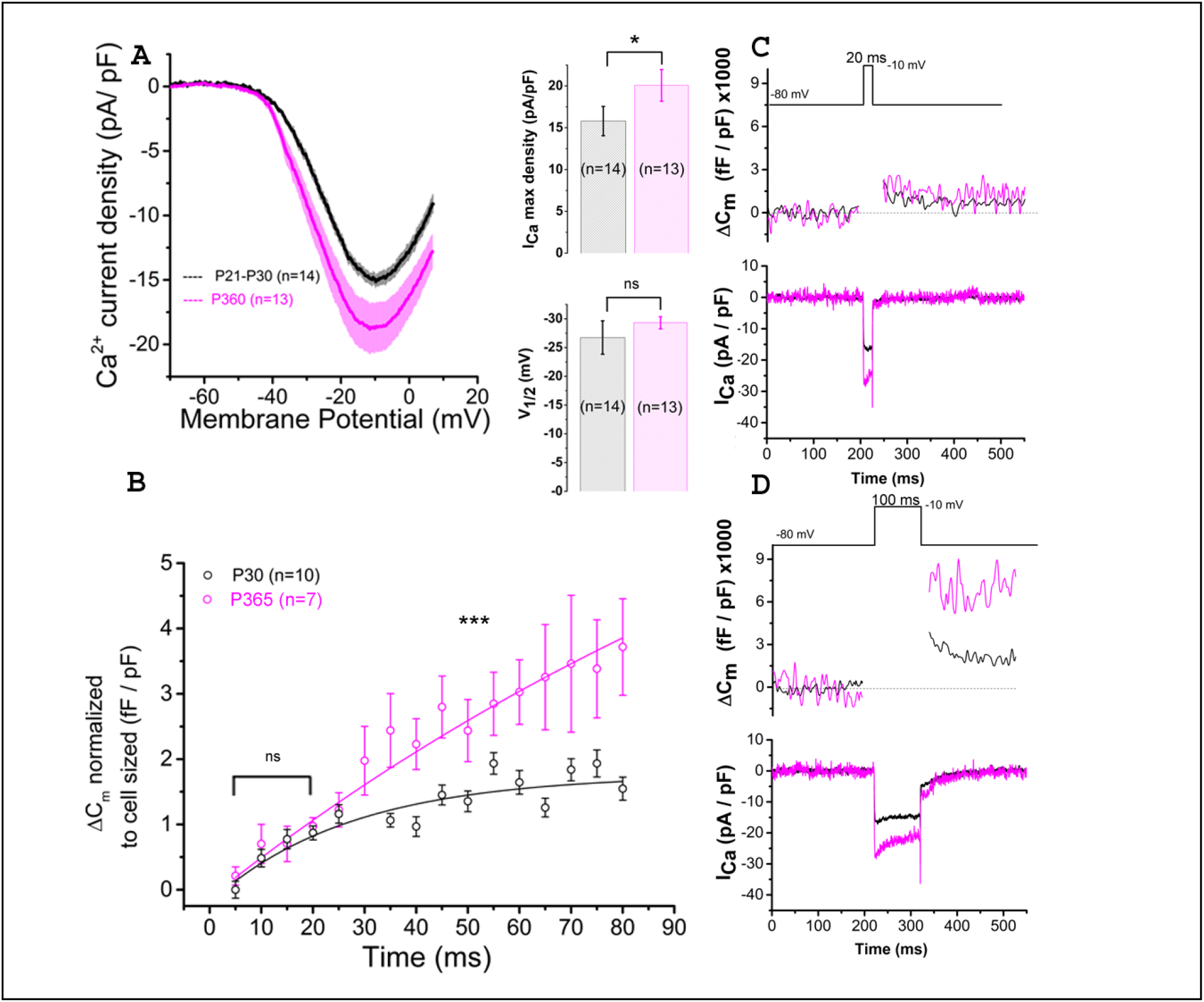
Ca^2+^ currents and exocytosis are exacerbated in IHCs from aged C57BL6/J mice. (**A**) Ca^2+^ current density of old P365 IHCs (pink) was larger as compared to young mature P21 IHCs (black). IHCs were subjected to a voltage ramp stimulation from -90 mV to +30 mV in 120 ms, giving a slope change in voltage of 1mV/ms. Comparative ICa maximum density measured at -10 mV indicated a significant increase in P360 IHCs (unpaired two-sample t-test, p=0.03) while the half-max voltage activation (V1/2) of the Ca^2+^ currents remained unchanged (unpaired t-test, p = 0.06). (**B**) Kinetics of exocytosis were evoked by a voltage-steps from -80 mV to -10 mV with increasing duration from 5 ms to 80 ms. Note that brief stimulations below 25 ms, a time-frame addressing the release of the Readily Releasable Pool of vesicles, the exocytotic response (normalized to cell size) was similar in P30 and P360 IHCs (p = 0.2; two-way ANOVA). Longer stimulations, above 30 ms, which probed the activation of the secondarily releasable pool of vesicles, showed a significant increase in exocytosis (two-way ANOVA; p = 2.3 E-8), indicating that old IHCs were more likely to sustain high rates of vesicular release. (**C-D**) Representative examples of ΔCm and ICa2+ recordings from P30 and P365 IHCs during a 20 ms and a 100 ms voltage-step stimulation from -80 mV to -10 mV. The number of IHCs, indicated in graph, were obtained from 5 mice at P365 and 6 mice at P30.

Patch-clamp measurement of membrane capacitance upon depolarizing voltage-steps from -80 to -10 mV with increasing time duration allowed to compare the kinetics of vesicle exocytosis in young and old IHCs. For brief stimulations below 25 ms, a time-frame addressing the release of the Readily Releasable Pool (RRP) of vesicles, the exocytosis response (normalized to cell size) was similar in P30 and P365 IHCs (Fig. 7B-C). However, longer depolarization above 25 ms revealed a more sustained exocytotic response in old IHCs, suggesting the mobilization of a more efficient secondarily releasable pool (SRP), i.e. a more efficient vesicle recruitment in these cells. To investigate whether this increased SRP exocytosis was due to higher Ca^2+^ microdomains at the ribbon active zones, we used the high affinity Ca^2+^-indicator Rhod-2 (K_d_ = 570 nM) which is known to allow a good monitoring of low Ca^2+^ concentrations ranging from 20 nM to 20 µM (Del Nido et al., 1998). We choose to use Rhod-2 rather than a low affinity Ca^2+^ dye such as OGB-5N with a Kd at 20 µM (Vincent et al., 2014) because we wanted here to primarily probe the amplitude of the Ca^2+^ microdomains, where the changes in Ca^2+^ concentrations are due to Ca^2+^ diffusion and are lower than at the nanodomain range near the Ca^2+^ channels. Ca^2+^ microdomains allow vesicle replenishment, a process occurring at the µm scale away from the ribbon and their associated Ca^2+^ channels (Spassova et al., 2004; Johnson et al., 2005; Graydon et al., 2011). We found that, upon 5 consecutive long 100 ms voltage-step stimulations (from -80 mV to -10 mV), IHCs displayed brighter transient Ca^2+^ hot spots (measured in 1µm^2^ areas) at their basal synaptic area in P365 mice as compared to young P30 IHCs (Fig. 8A-B). This indicated the expression of synaptic Ca^2+^ microdomains with larger amplitude in old P365 IHCs, a response likely due a larger Ca^2+^ channel density at their ribbon active zones (Fig. 7A). Simultaneously, in the same cells, the application of the 5 consecutive 100 ms stimulations, generated a facilitated vesicular recruitment in old P365 IHCs (Fig.8D). Also as a consequence of increased Ca^2+^ microdomains in old IHCs, the simultaneous recording of Ca^2+^ currents showed an enhanced time-inactivation, indicative of an increased Ca^2+^-dependent inactivation due to higher intracellular Ca^2+^ concentration (Fig. 8C).

**Figure 8.**
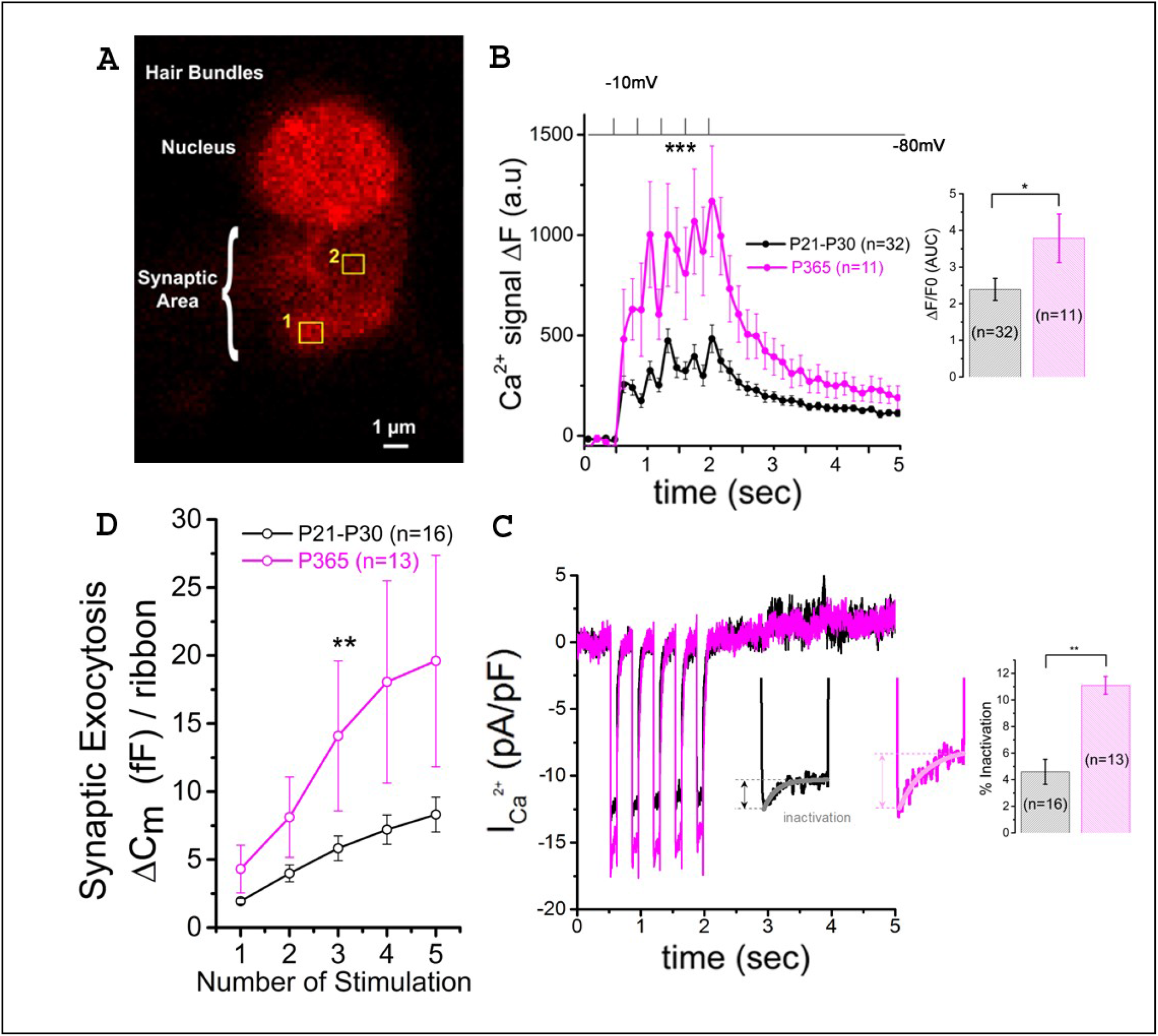
Larger synaptic Ca^2+^ signaling and increased sustained exocytosis in aged IHCs. (**A**) Example of confocal Ca^2+^ imaging, using the intracellular fluorescent Ca^2+^ indicator Rhod2, in an IHC. The cell was voltage-clamped at -80 mV and stimulated by 5 (100 ms) consecutive voltage-step to -10 mV. The photograph show two examples of square zones (1µm^2^) below the cell nucleus where voltage-induced calcium increase was measured. (**B**) Comparative mean amplitude of transient fluorescent hot spots showed higher Ca2+ responses at the presynaptic active zones of old P365 IHCs (pink) as compared to young mature P30 IHCs (black). Hair cells were depolarized by 5 five consecutive 100 ms depolarizing steps from -80 mV to -10 mV. The values n in the graph indicated the number of active zones analyzed for young (6 IHCs from 6 mice) and old mice (5 IHCs from 5 mice), respectively. The peak of the 5 calcium responses (ΔF=F-F0 with F0 as the fluorescent base line value averaged 0.5 s before stimulation) were significantly larger in old IHCs (two-way ANOVA (p = 8.8 E-5). The histogram compared the overall fluorescent response ΔF/F0 occurring during the 5 stimulations (0.5 to 2.5 s). The integral of the Ca^2+^ response (Area Under the Curve, AUC) was significantly larger in old P365 IHCs (unpaired t-test, p = 0.03). (**C**) Comparative mean of calcium current recorded at the same time during the 5 stimulations shown in B. Note that the inactivating part of the calcium current is increased in P365 IHCs. Asterisk indicate statistical analysis with unpaired t-test. (**D**) The concomitant exocytosis was also measured during the 5 stimulations shown in B Exocytosis measured as the variation of membrane capacitance was normalized for each IHC to their ribbon number or cell size; both normalization giving similar results). The exocytotic responses were larger in old P365 IHCs. The asterisks indicated statistical significant difference with p<0.05 (two-way ANOVA; p = 0.0015).

### Aging promoted autophagy in IHCs through the recruitment of LC3B

The size reduction of the old IHCs (Fig.6) suggested that the degenerative process of the ribbon synapses was associated with a concomitant reduction of the cell surface membrane of IHCs during aging, suggesting a largely augmented uptake of plasma membrane through endocytosis. Like other type of cells, such as neurons or retinal cells, auditory hair cells utilize autophagic pathways to maintain cell homeostasis during excessive ROS production (Defourny et al., 2019; Fu et al., 2018). Autophagy is a degradation process where cytoplasmic and plasma membrane components are engulfed by vesicles called autophagosomes. To determine whether this process could explain the IHC size reduction with aging, we investigated the recruitment of LC3B, a structural protein of autophagosomal membranes. We assessed and compared the autophagosome numbers and surface area in young P30 and old P365 IHCs by quantifying LC3B puncta numbers/cell by using confocal immunofluoresence. We found that the number and overall surface area of LC3-positive structures in IHCs largely increased with aging (Fig.9), indicating an increased autophagosome formation. Similar investigations in P365 CBA/J mice, a strain which does not display early age-related hearing loss (Fig. 1), showed a weak expression of LC3B in IHCs (Fig. 9).

**Figure 9.**
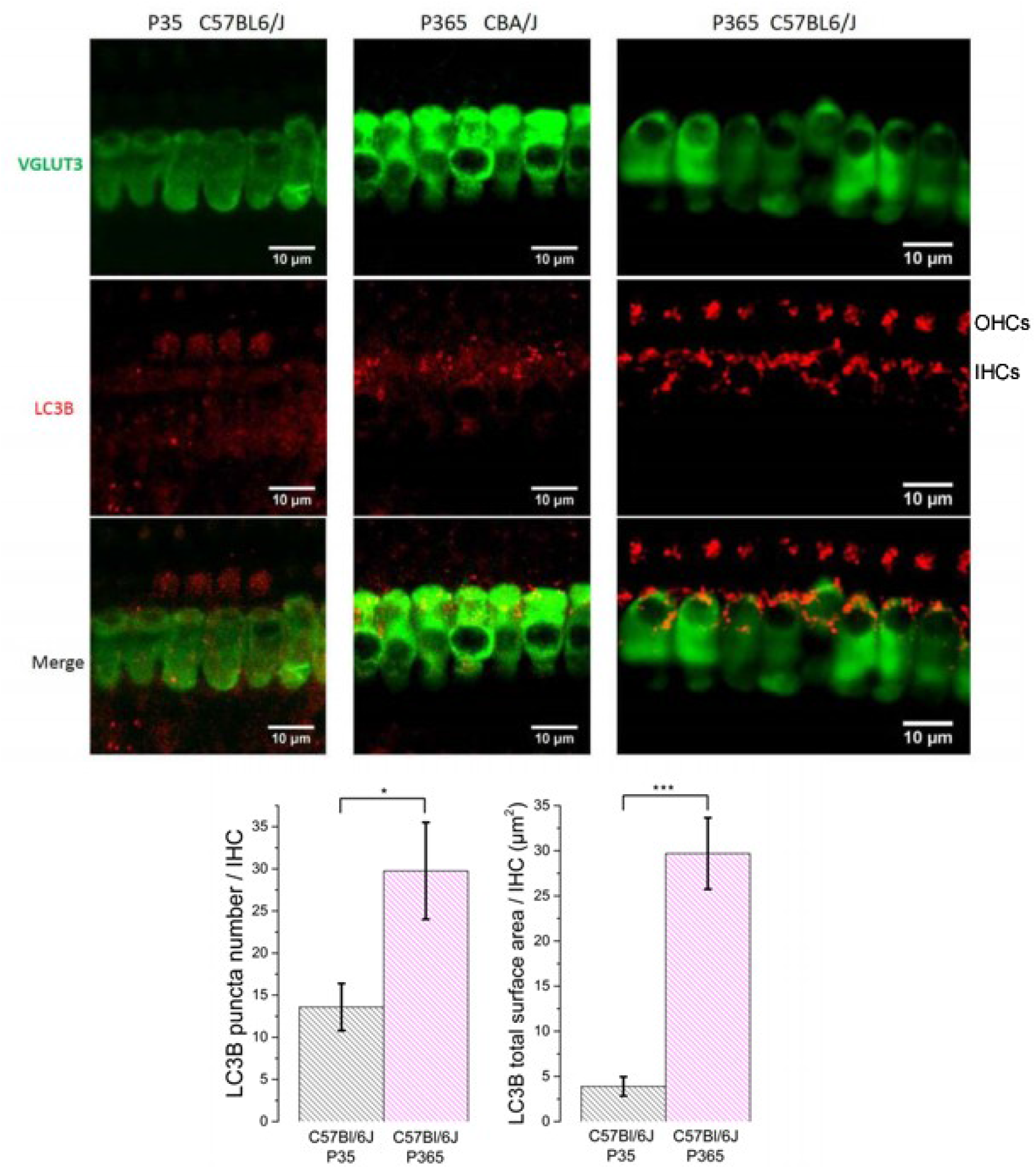
LC3-positive structures are prominent in aging IHCs. Representative confocal immunofluorescence images showing LC3B staining in IHCs from young P30 C57BL/6J (left), old P365 CBA/J (middle) and old P365 C57BL/6J (right) mice.The histograms below show the quantification of the LC3B puncta average number and average mean surface area per IHC. Images were obtained from processing confocal stack images (22 of 0.3 µm thickness) using ImageJ quantification tool (3D-object counter). Quantification was performed from 25 to 30 IHCs for each condition in the mid-cochlea (encoding region 8-16 kHz; 3 mice). Asterisks indicate statistical significance p < 0.05 (unpaired t test).

### Clustering and voltage-dependent activation of BK channels are affected aging in IHCs

There is increasing evidence that oxydative damage due to an increase in ROS levels plays a crucial role in age-related degeneration of the synaptic ribbon synapses (Fu et al., 2018; Rousset et al., 2020). BK (K_Ca1.1_) channels are known to be highly sensitive to oxydative stress and their activity is a good indicator of intracellular ROS production (Sahoo et al., 2014; Hermann et al., 2015). Interestingly, a decrease in BK channel expression in IHCs has been associated to noise-related ROS and peroxysome pathology (Delmaghani et al., 2015) but never yet described in IHCs during age-relate hearing loss. It is worth recalling that BK channels allows fast repolarization of IHCs and are essential for phase-locking of the auditory afferent neurons (Kros et al., 1998; Skinner et al., 2003; Oliver et al., 2006; Kurt et al., 2012). We here investigated, using immunofluorescent confocal microscopy, the comparative level expression of BK channels in IHCs from young (P30) and old P365 C57BL/6J mice. In young mature IHCs, BK channels are known to be organized in a dozen of clusters at the neck of the IHCs below the cuticular plate (Pyott et al., 2004; Hafidi et al., 2005). We used 3-D reconstructions from stacks of confocal images of immunostained organs of Corti (mid-cochlear frequency region 8-16 kHz) to determine the mean total surface area of the BK clusters per IHCs (Fig.10A-C). The results indicated that the membrane surface covered by BK channel clusters was reduced almost three fold in P365 IHCs as compared to P30 IHCs.

**Figure 10.**
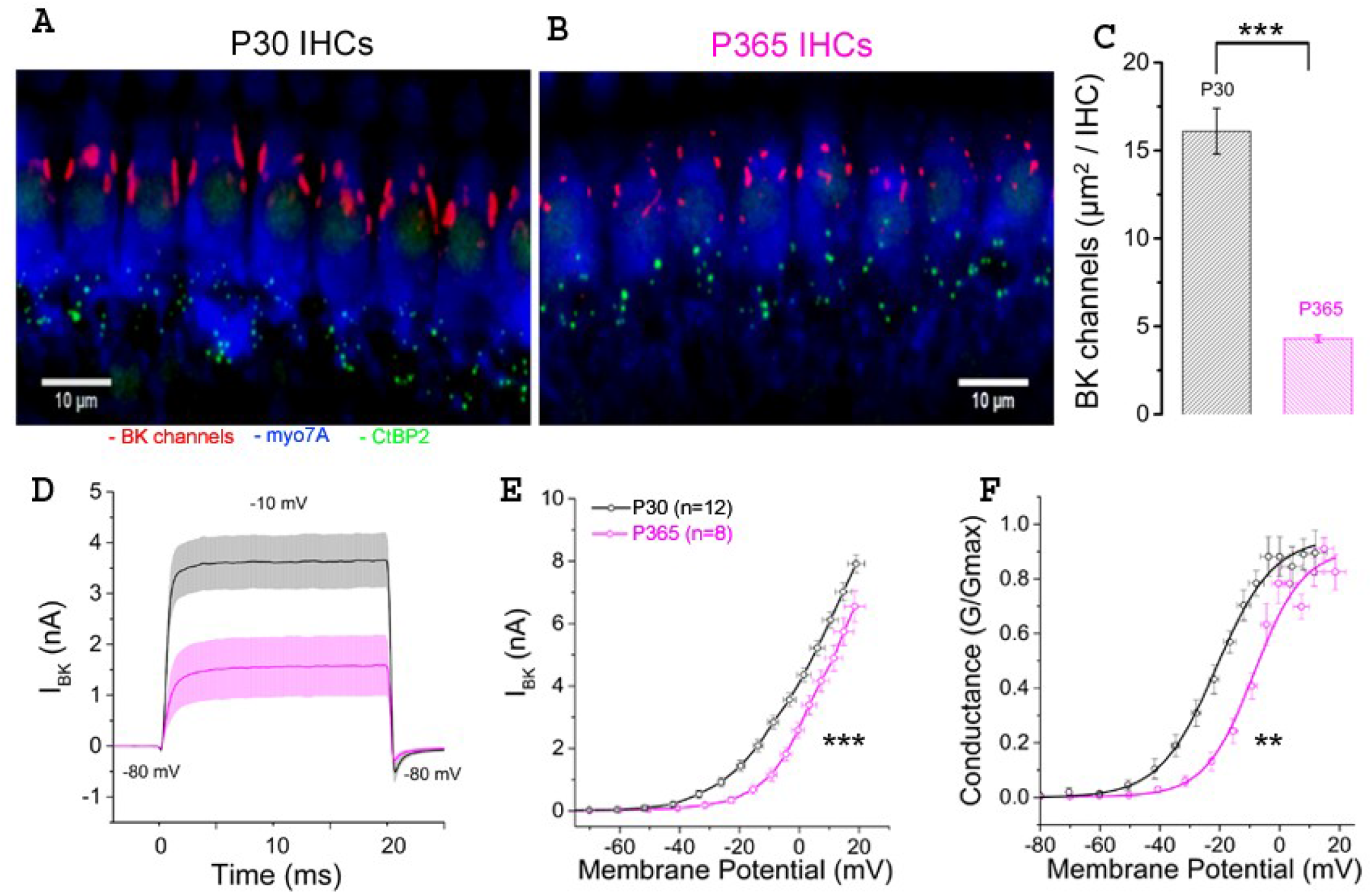
Immunofluorescent confocal imaging of BK channels in aging IHCs. (**A**) Numerous large BK channel clusters (red) are localized extrasynaptically below the cuticular plate of IHCs in young mature P30 C57BL6/J mice. (**B**) Note the strong reduction in size of the BK channels clusters in IHCs of old P365 C57BL6/J mice. (**C**) Histogram comparing the total surface area covered by the BK channel clusters per IHC in the mid-turn cochlea (region encoding 8-16 kHz). The total number of IHCs analyzed was 58 and 25 from P30 (n=3) and P365 (n=3) mice, respectively. Asterisks indicate statistical significance with p<0.05 (p = 1.12 E-7; unpaired t test). (**D**) Comparative fast outward potassium currents (IKf), carried by BK channels, recorded from P30 (black, n=12) and P360 (pink, n=8) IHCs when using a brief depolarization step of 20 ms from -80 to -10 mV. (**E**) Comparative BK current-voltage curves (n = 12 and 8 IHCs respectively in P30 and P360 mice, from 3 mice at each age). Significantly different two-way ANOVA, p = 2.7 E-8. IKf was measured 2.5 ms after the onset of the voltage-step stimulation. (**F**) Comparative BK conductance fitted with a Boltzmann curve indicated a V1/2 of -20.5 ± 5.7mV in P30 IHCs and of -13.7 ± 1.4 mV in P365 IHCs, respectively (unpaired t-test, p = 0.009). Note that voltage error across series resistance (Rs = 5 ± 2 MΩ) and leak current were compensated for each IHC. All values are expressed as means ± SEM.

**Figure 11.**
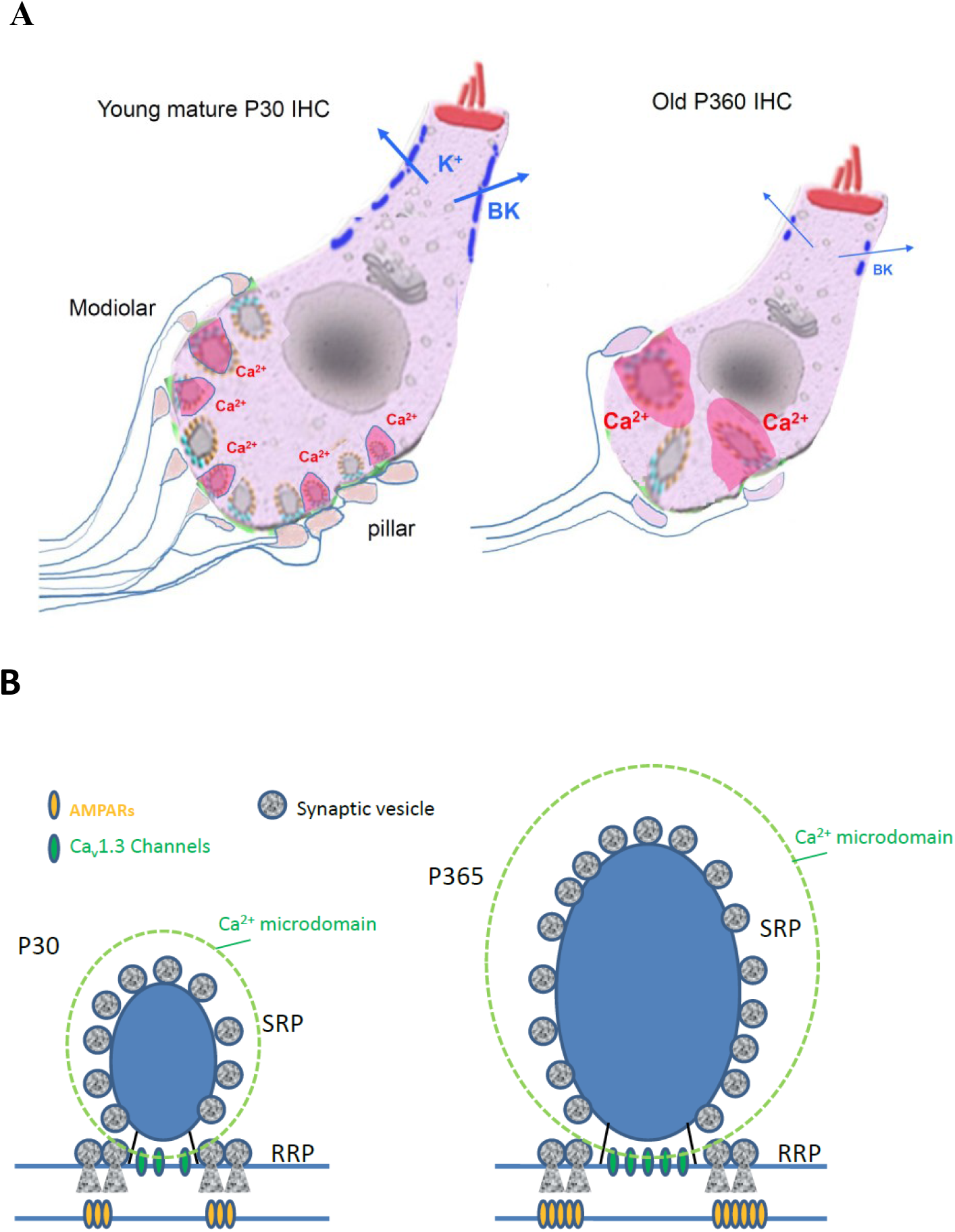
Synaptic IHC degeneration was associated with a strong potentiation of the remaining ribbon synapses in aging C57BL/6J mice. (**A**) Age-related hearing loss in P360 mice was associated with IHC shrinkage (cell size reduction), an important loss of afferent ribbon synapses (>50%), a drastic enlargement of the surviving synaptic ribbons (both at pillar and modiolar side), an increase in Ca^2+^ signaling (larger Ca^2+^ microdomain amplitude as indicated by the larger pink domains around the ribbons) and a large decrease in the fast repolarizing BK channel clusters (blue bars). (**B**) Hypothetical view of a young and old IHC ribbon synapse.

To further characterize the changes in BK channel function with aging, we performed whole cell patch-clamp recordings in IHCs from P30 and P365 mice. Since BK currents in IHCs have the peculiar property to be extremely fast activating currents (also termed IK_f_; Kros et al., 1998), these currents were recorded during brief 20 ms depolarizing steps and measured 2.5 ms after their onset (Fig.10D-F). BK currents in IHCs from young mature have also the particularity to activate at extremely negative membrane potential (near -60 mV) (Lingle et al., 2019). While the whole-cell conductance of BK channels from P365 IHCs was not significantly different from P30 IHCs (0.26 .± 0.04 nS and 0.23 ±.0.03 nS, respectively), their voltage-dependant activation showed a large positive shift of 7 mV (Fig.10F). Interestingly, the decrease in size of the BK patches and the positive voltage-activation shift in old IHCs somewhat resembled the phenotype observed in mice KO for LRRC52, a γ BK regulatory subunits (Lang et al., 2019; Lingle et al., 2019), suggesting that the BK complex formed with this γ subunit could be one the target of the degenerative oxydative process in aging C57BL56/J mice IHCs.

## DISCUSSION

### Larger synaptic ribbons associated with increased Ca^2+^ signals

Ribbons are presynaptic organelles tethering glutamate-filled vesicles at the hair cell active zone (AZ). These intracellular structures, composed of the protein RIBEYE, are anchored by the scaffold protein bassoon to the AZ plasma membrane, adjacent to Ca^2+^ Ca_V_1.3 channels (Brandt et al, 2003; Graydon et al., 2011; Jing et al., 2013; Vincent et al., 2017). Ribbons are essential for high rate of transmitter release and for high temporal precision of neuronal encoding of sound at the auditory afferent nerve fibers (Jean et al., 2018; Becker et al., 2018). Each auditory IHC have 10 to 20 synaptic ribbons depending of their cochlear frequency location (Meyer et al., 2009). Ribbons are heterogeneous in size and each ribbon makes a single synaptic contact with one afferent nerve fiber. Remarkably, fibers associated to large ribbons display low spontaneous rate and high-threshold have the tendency to be distributed at the modiolar side of IHCs (Liberman et al., 2011). Nerve fibers associated with smaller ribbons display high spontaneous rate and low-threshold activation, are mainly located toward the pillar side of IHCs. Interestingly, we found here that old C57BL6/J IHCs preferentially loose modiolar ribbons (known to be associated with high threshold low spontaneous fibers), therefore mimicking what is happening during noise-induced hearing loss (Fernandez et al., 2015; Liberman and Kujawa 2017; Hickman et al., 2020). Heterogeneity in the voltage-dependence of Ca^2+^ channels and release site coupling has been recently proposed to contribute to the firing fiber specificity, with modiolar ribbons being activated at more negative potentials (Özçete and Moser 2021). In our study, we did not find evidence for a voltage-shift in the voltage-dependence of Ca^2+^ channels with aging but we did observe an increase in the density of Ca^2+^ currents. The increase in Ca^2+^ entry was further confirmed by the stronger time-inactivation of the Ca^2+^ currents (Grant and Fuchs, 2008; Vincent et al., 2017).

One of the most striking observations was that the IHC synaptic ribbons underwent a nearly threefold increase in size with aging, confirming a certain degree of plasticity of these synaptic structures. The enlargement of the presynaptic ribbons was in good agreement with a previous ultrastructural study in aging C57BL/6J mice by Stamataki et al. (2006) and the recent study of Jeng et al., (2020). Interestingly, the size of the young mature IHC synaptic ribbons is known to correlate positively with the AZ Ca^2+^ microdomain amplitude, as well as the number and gating of the Ca_v_1.3 Ca^2+^ channels (Frank et al., 2009; Neef et al., 2018). Also, hair cells of transgenic zebrafish with enlarged ribbons display larger ribbon-localized Ca^2+^ currents (Sheets et al., 2017). In agreement with these previous studies, we found here, in IHCs from aging C57BL/6J mice, that the remaining larger ribbons were associated with larger Ca^2+^ microdomain amplitude, and larger density of calcium currents, reinforcing the idea that Ca^2+^ ions regulate the size of the synaptic ribbons. In this context, It is worth recalling that mitochondrial-Ca^2+^ uptake has been proposed to lower cellular NAD^+^/NADH redox and regulates ribbon size, possibly by acting through the NAD^+^ binding site of the B domain of RIBEYE (Wong et al., 2019). In vitro work has shown that NAD^+^ and NADH can promote interactions between ribeye domains (Magupalli et al., 2008) and the size of the ribbons would be then sensitive to the subcellular synaptic NAD^+^/NADH redox level (Wong et al., 2019), suggesting that the increase in size of the IHCs ribbons from aged C57BL/6J mice also results from NAD^+^/NADH homeostasis disruption.

### Increased capacity for sustained IHC synaptic exocytosis: Implication for the discharge pattern of the auditory nerve firing

As a consequence of larger Ca^2+^ microdomains, we found that IHCs from old C57BL/6J mice had larger and more sustained exocytotic responses. These larger release responses could also well be explained by a higher synaptic vesicle density at the active zone of the IHC ribbon synapses as suggested by the EM study of Stamataki et al. (2006). Surprisingly, the recent study of Jeng et al., (2020), which did not perform Ca^2+^ imaging, concluded that the size and kinetics of Ca^2+^-dependent exocytosis in IHCs were unaffected in aging mice but this study did not normalize the IHC exocytotic responses to their cell size or to their ribbon synapse number, and therefore may have overlooked the differential phenotype.

What would be the consequences on the discharge pattern of the remaining afferent fibers of a more sustained release capacity in old ribbon synapses ? Fast adaptation in the discharge rate of the afferent auditory nerve fibers is thought to be one of their essential property for emphasizing the timing information of sound stimuli. This adaption takes the form of a fast increase in spike rate at stimulus onset that rapidly declines in a double exponential decay to a lower steady-state rate after a few tens of milliseconds (Taberner and Liberman 2005). The origin of this fast adaptation has been proposed to arise first from H^+^ release block of Ca^2+^ channels and second to rapid depletion of synaptic vesicle release (Vincent et al., 2017). We predict that the increased capacity to sustain SRP exocytosis in old IHCs with larger ribbons would have the tendency to decrease the level of firing adaptation of the auditory fibers and therefore explain the altered recovery from short-term adaptation in old C57BL/6J mice (Walton et al., 1995).

Furthermore, we confirmed here the hyperacusis-like effect of the startle responses in old C57BL/6J mice (Ison et al., 2007). This increased response at low sound intensity could be explained by a release potentiation at the IHC ribbon synapses due to the presynaptic higher Ca^2+^ channel density associated with a larger ribbon coupled with a larger vesicular SRP, and increased expression of post-synaptic AMPARs. All these features would make the auditory ribbon synapses hypersensitive and underlie the hyperacusis-like reflex in old C57BL/6J mice.

### IHC shrinkage and disruption of BK channel clusters

Another novel observation of our study was the drastic reduction in cell size of IHCs from aging C57BL/6J mice. Interestingly, tuberous sclerosis rapamycin-sensitive complex 1 (mTORC1) has been shown to be overactivated and promote oxidative damages to synapse ribbon loss in aging C57BL/6 mice (Fu et al., 2018). We found that aged P365 IHCs had an increase expression of the autophagy marker LC3B. The activation of the mTOR pathway with aging could therefore trigger the cell size reduction of IHCs, by activating the autophagy pathway, as proposed in many other cell types (Fingar et al., 2002; Fumarola et al., 2005). In addition, our study described another novel remarkable feature of aging IHCs, a drastic disruption of the BK channel clusters, suggesting that these potassium channels are also a good indicator of the redox state and autophagy in IHCs. The expression and distribution of BK channels could be regulated through the rapamycin insensitive mTOR complex 2 (mTORC2) as shown in the kidney glomerulus podocytes (Wang et al., 2019). Interestingly, the disruption of the BK channel clusters in aging IHCs was associated with a large positive shift in their voltage-activation curve, suggesting that the fast repolarizing property mediated by BK channels which likely contributes to decrease transmitter release (Skinner et al., 2003), would be largely hampered during sound stimuli in aging mice, an additional factor that may also contribute to hyperacusis.

## CONCLUSIONS

In conclusion, our study unraveled several novel functional and structural features of aging in IHC ribbon synapses from C57BL/6J mice (Fig.10): 1) a large reduction in IHC cell size, 2) a drastic enlargement of the presynaptic ribbons and postsynaptic AMPAR clusters 3) an increased presynaptic Ca^2+^ signaling associated with a stronger sustained exocytotic response 4) a disruption of BK channel clusters and a drastic shift in their voltage-dependence decreasing their potential negative feedback on neurotransmission. Overall, these results suggested that aging IHCs ribbon synapses can undergo important structural and functional plasticity that lead to synaptic potentiation that could explain the paradoxical hyperacusis-like exaggeration of the acoustic startlle reflex observed in C57BL/6 mice.

## MATERIALS and METHODS

### Mice

Inbred C57BL/6J mice, which are homozygous for the recessive cadherin23 mutation *Cdh23*^*753A*^ (also known as Ah1) were used in the present study. Hearing thresholds and IHC ribbon counts in mice of either sex were performed at postnatal ages: P15, P30, P60, P180 and P365. CBA/J mice (from Charles River) were also tested as controls at P50, P250 and P365. Animal care and all procedures were approved by the institutional care and use committee of the University of Bordeaux and the French Ministry of Agriculture (agreement C33-063-075).

### Assessment of hearing thresholds (ABRs)

To record ABRs (Auditory Brainstem Responses, which represent the sound-evoked synchronous firing of the auditory cochlear nerve fibers and the activation of the subsequent central auditory relays), mice were anesthetized with intraperitoneal injection of xylazine (6 mg/ml, Rompun Bayer) and ketamine (80 mg/ml, Virbac) mixture diluted in physiological saline. The mouse was placed in a closed acoustic chamber and its body temperature was kept constant at 37°C during ABRs recording. For sound stimulus generation and data acquisition, we used a TDT RZ6/BioSigRZ system (Tucker-Davis Technologies). Click-based ABR signals were averaged after the presentation of a series of 512 stimulations. Thresholds were defined as the lowest stimulus for recognizable wave-I and II. The amplitude of ABR wave-I was estimated by measuring the voltage difference between the positive (P1) and negative (N1) peak of wave-I. Sound intensities of 10 to 90 dB SPL in 10 dB step, were tested.

Distortion product otoacoustic emissions (DPOAEs), which originate from the electromechanical activity of the outer hair cells (OHCs), were tested by using two simultaneous continuous pure tones with frequency ratio of 1.2 (f1 = 12.73 kHz and f2 = 15,26 kHz). DPOAEs were collected with the TDT RZ6/BioSigRZ system designed to measure the level of the “cubic difference tone” 2f1-f2.

### Acoustic Startle Reflex (ASRs) measurements

Testing was conducted in four acoustic startle chambers for mice (SR-LAB, San Diego Instruments, San Diego, CA). Each chamber consisted of a nonrestrictive cylindrical enclosure attached horizontally on a mobile platform, which was in turn resting on a solid base inside a sound-attenuated isolation cubicle. A high-frequency loudspeaker mounted directly above the animal enclosure inside each cubicle produced a continuous background noise of 65 dBA and various acoustic stimuli in the form of white noise. Mechanical vibrations caused by the startle response of the mouse were converted into analog signals by a piezoelectric accelerometer attached to the platform. A total of 130 readings were taken at 0.5-ms intervals (i.e., spanning across 65 ms), starting at the onset of the startle stimulus. The average amplitude over the 65 ms was used to determine the stimulus reactivity. The sensitivity of the stabilimeter was routinely calibrated to ensure consistency between chambers and across sessions.

Acoustic startle reflex was assessed during a session lasting for approximately 30 minutes, in which the mice were presented with a series of discrete pulse-alone trials of different intensities and durations. Ten pulse intensities were used: 69, 73, 77, 81, 85, 90, 95, 100, 110, and 120 dBA lasting for either 20 or 40 ms (background noise level: 65 dBA). A session began when the animals were placed into the Plexiglas enclosure. The mice were acclimatized to the apparatus for 2 minutes before the first trial began. The first six trials consisted of six pulse-alone trials of 120 dBA, comprising three trials of each duration. These trials served to stabilize the animals’ startle response, and were analyzed separately. Subsequently, the animals were presented with five blocks of discrete test trials. Each block consisted of 20 pulse alone trials, one for each intensity and duration. All trials were presented in a pseudorandom order with an inter-trials interval of 14 seconds.

### Tissue preparation and imunocytochemistry

Mice were deeply anesthetized with isofluorane (Vetflurane, Virbac). The ramps of the cochlear apparatus were dissected and prepared as previously described by Vincent et al. (2014). Inner ears (cochleae) were fixed by incubation in 4% paraformaldehyde in phosphate-buffered saline (PBS), pH 7.4, at 4°C overnight and washed with cold phosphate buffered saline (PBS). They were then incubated for 2 days in PBS solution containing 10% EDTA pH 7.4 at 4°C. The middle part of the organ of Corti (area encoding between 8 and 16 kHz) was then dissected and the tectorial membrane removed. The tissue was first incubated with PBS containing 30% normal horse serum and triton X100 0.5 % for 1 hour at room temperature. Synaptic ribbons were labeled with anti-CtBP2 (1/200 Goat polyclonal, Santa Cruz, USA ; cat # SC-5966 or 1/200 Mouse IgG1 monoclonal antibody BD Biosciences, cat # 612044). Otoferlin was labeled with a mouse monoclonal antibody (1/200 AbCam, cat # ab53233). Autophagy was detected by labeling LC3B protein with anti Rabbit monoclonal antibody (1/200 LC3B(D11) XP, Cell Signaling Technology, cat # 3868). BK channels were labeled with a Rabbit polyclonal antibody (1/200, Alomone, Israel, cat #APC-021. Explants of Organ of Corti were incubated overnight at 4°C with primary antibodies. The following fluorescent secondary antibodies 1/500 were then used: anti-Goat Donkey polyclonal Fluoprobes 547H (Interchim, cat# FP-SB2110), anti-Mouse Donkey polyclonal Fluoprobes 647H (Interchim, cat# FP-SC4110), anti-Mouse Donkey polyclonal Alexa Fluor 488 (AbCam, cat# ab150109), anti-Mouse Goat polyclonal Alexa Fluor 546 (Molecular Probes, cat# A-11003), anti-Mouse Goat polyclonal Alexa Fluor 488 (Molecular Probes, cat# A-10680), anti-Rabbit Donkey polyclonal Fluoprobes 594 (Interchim, cat# FP-SD5110). Actin-F was also used to visualize hair cells (1/100, Phalloidin Fluoprobe 405, Interchim, Montlucon, France; cat # FP-CA9870). For Image Acquisition, organ of Corti samples were analyzed using a confocal laser scanning microscope Leica SP8 with a 63X oil immersion objective (NA = 1.4) and white light laser (470 to 670 nm) (Bordeaux Imaging Center). Phalloidin was imaged by using a diode laser at 405 nm also mounted on the microscope. For complete 3D-stack reconstruction of IHCs, 25 to 30 images (0.3 µm thickness) were acquired as previously described (Vincent et al, 2014). Ribbon volumes were calculated using the 3D-Objects Counter Plugin of ImageJ. Image resolution was 0.038 µm/pixel. As previously described by Hickman et al. (2020) and Jeng et al., (2020), the x,y,z coordinates of ribbons in each z-stack were transformed into a coordinate system based on the modiolar-pillar polarity of the IHCs. We determined the pillar versus modiolar distribution of the synaptic ribbons by using a *k*-means clustering method in the z-stack position of each ribbon, with arbitrarily choosing the number of cluster as 2 (OriginPro 9.1software, OriginLab, Northampton, USA). From these 2 clusters, we determine arbitrarily the border between the modiolar and pillar side in each IHC by taking the half-distance between the two nearest ribbon of each cluster.

### High resolution imaging of synaptic ribbons with confocal STED microscopy

For high-resolution imaging of IHC ribbons, we used a Leica DMI6000 TCS SP8 STED microscope with 93X glycerol immersion objective (NA 1.3) (Bordeaux Imaging Center). Mouse sensory organs were fixed and incubated with primary anti CtBP2 (1/100) antibodies in conditions similar to those described above for confocal microscopy. Secondary antibodies Goat anti-IgG1 Mouse polyclonal Alexa Fluor 647 (Jackson, cat# 115-605-205) were used to label CtBP2 primary antibodies. The pulsed STED depletion laser (pulsed laser IR, Spectra Physics Mai Tai) was set at 775 nm. Stack images were acquired with the following parameters: scan rate 200 Hz, 775 nm depletion laser power between 8 and 15%, scans frames accumulated per *X-Y* section 8 times, z-step size 0.25 µm, number of plans between 8 and 10, pixel size 20 nm giving an *X-Y* image size of 21 × 5 μm (1024 × 256 pixels).

### Counting spiral ganglion neurons

The afferent nerve fibers of the auditory nerve were fluorescently labeled using a retrograde track tracing technique with dextran amines (Fritzsch et al., 2016). Cochleae were extracted from temporal bone of freshly euthanised mice and rapidly immerged intact into ice-cold artificial perilymph (NaCl 135; KCl 5.8; CaCl_2_ 1.3; MgCl_2_ 0.9; NaH_2_PO_4_ 0.7; Glucose 5.6; Na pyruvate 2; HEPES 10, pH 7.4, 305 mOsm). A small crystal of tetramethylrhodamine dextran amines 3000 (Life technologies, NY) was placed for 15-20 min at the edge of the auditory nerve coming out at the internal auditory canal of the intact cochlea and rinsed with artificial perilymph. The intracochlear sensory tissues were then fixed by gentle perfusion of a 4% PFA solution for 30 min and rinsed with PBS. They were then decalcified for 2 days in PBS solution containing 10% EDTA pH 7.4 at 4°C. The cochleae were then sliced with a razor blade along the longitudinal cochlear axes and the slices were then mounted on a glass-slide for fluorescent confocal microscopy observation. The quantification of the spiral ganglion neurons per surface area of the neural ganglion were made in the middle part of the cochlea in a region encoding 8-16 kHz, by using the 3D object counter application of ImageJ software.

### Ca^2+^ currents and whole-cell membrane capacitance measurement in IHCs

Measurements of resting membrane capacitance of IHCs were performed at P30 and P365 mice in the 20-40% normalized distance from the apex, an area coding for frequencies ranging from 8 to 16 kHz, by using an EPC10 amplifier controlled by Patchmaster pulse software (HEKA Elektronik, Germany), as previously described in detail (Vincent et al., 2014, 2015). The organ of Corti was incubated in an extracellular perilymph-like solution containing NaCl 135; KCl 5.8; CaCl_2_ 5; MgCl_2_ 0.9; NaH_2_PO_4_ 0.7; Glucose 5.6; Na pyruvate 2; HEPES 10, pH 7.4, 305 mOsm. This extracellular solution was complemented with 0.25 µM of apamin (Latoxan; cat # L8407) and 1 µM of XE-991 (Tocris Bioscience; cat # 2000) to block SK channels and KCNQ4 channels, respectively. The external Ca^2+^ concentration was increased from 1.3 to 5 mM to enhance the amplitude of Ca^2+^ currents to levels nearby body temperature. All experiments were performed at room temperature (22°C–24°C).

Patch pipettes were pulled with a micropipette Puller P-97 Flaming/Brown (Sutter Instrument, Novato, CA, USA) and fire-polished with a Micro forge MF-830, (Narishige, Japan) to obtain a resistance range from 3 to 5 MΩ. Patch pipettes were filled with a cesium-based intracellular solution containing (in mM): CsCl 145; MgCl_2_ 1; HEPES 5; EGTA 1; TEA 20, ATP 2, GTP 0.3, pH 7.2, 300 mOsm. Measurements of the resting membrane capacitance (cell size) of IHCs were obtained in whole-cell voltage-clamp configuration at -70 mV and after 2 min equilibrium of the internal patch-pipette recording solution with the IHC cytosol.

### BK currents

Whole-cell recordings of BK currents were recorded in P30 and P365 IHCs from ex-vivo whole-mount preparation (Skinner et al., 2003) as described above for Ca^2+^ currents. Recording pipettes were filled with a KCl-based intracellular solution containing in mM: 158 KCl, 2 MgCl2, 1.1 EGTA, 5 HEPES, and 3.05 KOH, pH 7.20.

### Intracellular Ca^2+^ imaging

Fluorescent Ca^2+^ signals were measured with a C2 confocal system and NIS-element imaging software (Nikon) coupled to the FN1 upright microscope as previously described (Vincent et al., 2014 ; 2017). For Ca^2+^ imaging in IHCs, in which membrane capacitance and Ca^2+^ currents were simultaneously recorded, patch pipettes were filled with the same cesium-base solution described above supplemented with 50 µM of the red fluorescent indicator Rhod2 (R14220, Invitrogen™, Thermofisher Scientific). The Ca^2+^ dye was excited with a 543 nm Helium Neon Laser system (Melles Griot 05-LGR-193-381) coupled to a Nikon C2-confocal FN1-upright microscope and emission fluorescence at 552-617 nm was recorded at 18 images/s (resolution 0.4 µm/pixel). Using the Nikon imaging system NIS-Elements, IHC active zones were first individually identified by their fluorescent transient responses, following a 100 ms stimulation (from -80 to -10 mV) and then the focal plane (with pinhole 1.1 µm) was adjusted in the z dimension to give the brightest value for each active zone. Then, after a 1 min rest, each active zone was stimulated by a train of five consecutive 100 ms stimulation, each separated by 250 ms. Fluorescence emission in each active zone was subsequently measured with ImageJ software by drawing a region of interest of 1 μm^2^ at the synaptic zone (Fig.9B). Emission fluorescent signals were analyzed and normalized by the ratio Δ*F/F*_0_ ratio, where *F*_0_ was the fluorescence level before stimulation.

### Statistical analysis

Results were analyzed with OriginPro 9.1software (OriginLab, Northampton, USA). Results were analyzed with OriginPro 9.1software (OriginLab, Northampton, USA). Data were tested for normal distribution using the Shapiro-Wilk normality test, and parametric or nonparametric tests were applied accordingly. For statistical analyses with two data sets, two-tailed unpaired t-tests or two-sided Mann-Whitney tests were used. For comparisons of more than two data sets, one-way ANOVA or two-way ANOVA followed by Tukey multiple comparison test were used when data was normally distributed. If a data set was not normally distributed Kruskal-Wallis tests followed by Dunn multiple comparison tests, respectively, was used. When more than two data sets were compared and significant differences were found, reported p values correspond to the post-hoc multiple comparison test. Values of p < 0.05 were considered significant. Results are expressed as mean ± SEM.

## Funding & acknowledgments

This work was supported by a grant from the Fondation Pour l’Audition to DD (IDA CLINICAL TRANSLA - 010703E), and from the French Society of Audioprosthesis professionals, Entendre SAS (Saint Cyr l’Ecole 78210 France). Part of the content of the manuscript has previously appeared as a preprint online at bioRxiv https://doi.org/10.1101/2020.05.15.097550

